# Temporal modulation of tactile perception during balance control

**DOI:** 10.1101/2023.11.02.565260

**Authors:** Fabian Dominik Wachsmann, Katja Fiehler, Dimitris Voudouris

**Affiliations:** Experimental Psychology, Justus Liebig University Giessen, Otto-Behaghel-Str. 10F, 35394, Giessen, Germany

**Keywords:** perception & action, postural adjustments, tactile modulation, virtual reality

## Abstract

Somatosensory feedback is essential for motor control, yet sensations from one’s own movements are often suppressed. Despite extensive research on movement-induced tactile suppression, its mechanisms remain unclear. Most studies focus on simple upper-limb movements, leaving the generalization to other body parts unexplored. This study examines tactile processing on the lower limb during balance control by varying feedback processing demands. Participants experienced visual perturbations in a virtual room challenging their posture. Tactile sensitivity was assessed using vibrotactile stimuli to the lower leg at different times around the perturbation. We found that postural behavior is both predictively tuned before and reactively adjusted to expected perturbations. Our results provide evidence that tactile sensitivity changes according to feedback processing demands on the lower limb. Such dynamic sensory modulation could reflect the continuous up- and down-regulation of feedback signals to accomplish the task at hand.

## Introduction

Tactile sensitivity is reduced on a moving limb compared to the same but static limb. This is typically reflected in a lower perception of externally-generated probe stimuli delivered to a limb while being engaged in movement planning or execution, as shown for simple finger extensions, to reaching, grasping and even juggling (Buckingham et al., 2010; Chapman & Beauchamp, 2006; Juravle & Spence, 2011; Voss et al., 2008; Voudouris & Fiehler, 2017a). Such reduction in tactile sensitivity has been demonstrated both on the perceptual (Buckingham et al., 2010; Voss et al., 2008; Voudouris & Fiehler, 2017) and neural level (Parkinson et al., 2011; Seki & Fetz, 2012; Arikan et al., 2021). Although tactile suppression may be explained by peripheral mechanisms, such as reafferences that mask the relatively weak probe stimuli (Williams & Chapman, 2002; Chapman & Beauchamp, 2006; see also Broda et al., 2020), the prevailing idea is that sensorimotor predictions down-weight the incoming sensory feedback arising from the movement (Shergill et al., 2003; Voss et al., 2006).

On the perceptual level, tactile suppression can be modulated by various factors. For instance, when sensory feedback from the moving limb is highly predictable, tactile suppression on that limb increases (Chapman & Beauchamp, 2006; Fuehrer et al., 2022). On the other hand, tactile suppression is reduced when sensory feedback from the moving limb is needed to accomplish the task, e.g. when grasping objects of unknown properties or of low friction (Voudouris et al., 2019; Voudouris & Fiehler, 2022). Tactile suppression is also modulated throughout the execution of arm movements (Colino & Binsted, 2016; Juravle et al., 2018; Voudouris & Fiehler, 2021), showing reduced suppression at moments when sensory guidance of a reaching movement gains importance (Voudouris & Fiehler, 2021). All in all, these results suggest that tactile sensitivity on a body part depends on the dynamic interplay between central predictive and somatosensory feedback signals.

Despite the large body of research on this phenomenon, mechanisms underlying up- and down-regulation of tactile perception during movement are still unclear. Some studies found that somatosensory-evoked potentials related to tactile stimulation on the foot sole are decreased (Morita et al., 1998) or increased (Mouchnino et al., 2015) prior to voluntary foot flexion or step initiation, respectively. This apparent discrepancy in physiological signal processing may arise from the different need to use tactile feedback from the foot sole to accomplish the task: prior to initiating a step (Mouchnino et al., 2015), tactile sensitivity may improve as tactile feedback from the foot sole provides crucial information for balance control. In contrast, tactile feedback from the foot sole prior to an arbitrary flexion of the foot (e.g. Morita et al., 1998) may be less relevant for movement execution and thus it may be down-regulated. In the current study, we test the hypothesis that tactile sensitivity is modulated by the need to process tactile feedback from the lower leg that is involved in the retention of upright posture. We applied a common psychophysical approach allowing us to compare our results to previous work on tactile suppression during upper limb movements (cf., Fuehrer et al., 2022; Kilteni & Ehrsson, 2022; Thomas et al., 2021).

While continuous sensory integration is crucial for balance control (Peterka, 2002; Chiba et al., 2016), less is known about how tactile signals from the standing body are modulated by feedback processing demands. For instance, when balance is perturbed, integration of visual, vestibular, and somatosensory feedback signals is necessary to form a motor plan and elicit required adjustments. Tactile signals from the foot are shown to be important as they are integrated into the postural motor plan for the upcoming initiation of a step (Mouchnino & Blouin, 2013), whereas disrupting the quality of tactile information from the foot can result in poorer upright stance (Fiolkowski et al., 2002; Voudouris et al., 2013). Meanwhile, tactile information from the lower leg can be exploited to support body balance (Menz et al., 2006) and to obtain a better estimate of the current body configuration to facilitate postural adjustments. Based on these, if tactile processing is modulated at moments when somatosensory control of posture gains importance, we expect improved tactile sensations on the lower leg just before the onset of a postural adjustment to a perturbation.

To examine how tactile sensitivity on the lower leg is modulated by feedback processing demands, we asked participants to stand within a virtual room, facing a wall that would move towards them with low or high temporal uncertainty. This visual perturbation was introduced to challenge balance control (Chander et al., 2019) by allowing for merely reactive (high temporal uncertainty) or predictive (low temporal uncertainty) postural adjustments to the perturbations. To probe tactile sensitivity, we presented a brief tactile stimulus of varying intensity to the participant’s lower leg at various moments prior to and immediately after the onset of the visual perturbation. This approach is similar to examining tactile sensitivity during upper limb movements when the tactile probe is delivered around the area of the involved effector but not directly to the skin area involved in the manual interaction (e.g., fingertip during grasping; Chapman et al., 1987; Buckingham et al., 2010; Fuehrer et al., 2022). Here, we probed the skin over the calf muscle that is in close proximity to the foot sole (direct contact area), and can convey tactile signals that can improve body posture (Menz et al., 2006). It is important to note that the probing tactile stimuli are neither informative nor relevant for maintaining upright stance in our study.

To confirm that our manipulation had the expected effects on body posture, we examined whether postural sway would be reduced in anticipation to perturbations of low temporal uncertainty (cf. Eikema et al., 2013) and whether distinct body reactions would occur after the onset of either type of perturbation, always relative to quiet stance. We further expected altered tactile sensitivity during standing in perturbation compared to quiet stance. If feedback processing demands modulate tactile sensitivity on the lower limb, we expected increased tactile sensitivity at moments when postural control is deemed more important (i.e. around the time of a postural reaction). Finally, to explore if tactile sensitivity is modulated by temporal uncertainty and related postural demands, we examined if it differs around the onset of the two types of perturbations (high vs low temporal uncertainty).. In a second experiment, we explored whether the requirement to keep upright imposes any further changes in tactile sensitivity on the lower limb. To this end, we tested whether tactile sensitivity changes between standing and sitting.

## Results

### Experiment 1

#### Kinematics

As expected, anticipatory postural control was generally evident prior to the low uncertainty perturbation compared to the respective baseline. Specifically, the maximal displacement of the centre of pressure (COP) was smaller than that during the respective baseline (*t*_19_ = 2.114, *p* = 0.024, *η^2^* = 0.053; Figure 1A), in line with our hypothesis based on previous work (Eikema et al., 2013). The effect on the head’s maximal displacement was not as systematic (*t*_18_ = 1.420*, p =* 0.086*, η^2^ =* 0.026; Figure 1B). Meanwhile, there was no evidence of anticipatory behavior prior to the high uncertainty perturbation, as reflected both in the COP and the head maximal displacements (both *t* <= 0.202, both *p >*= 0.842, both *η^2^* <= 0.001). These findings confirm that participants considered the underlying dynamics of the perturbations and predictively tuned their posture whenever possible.

**Figure 1:**
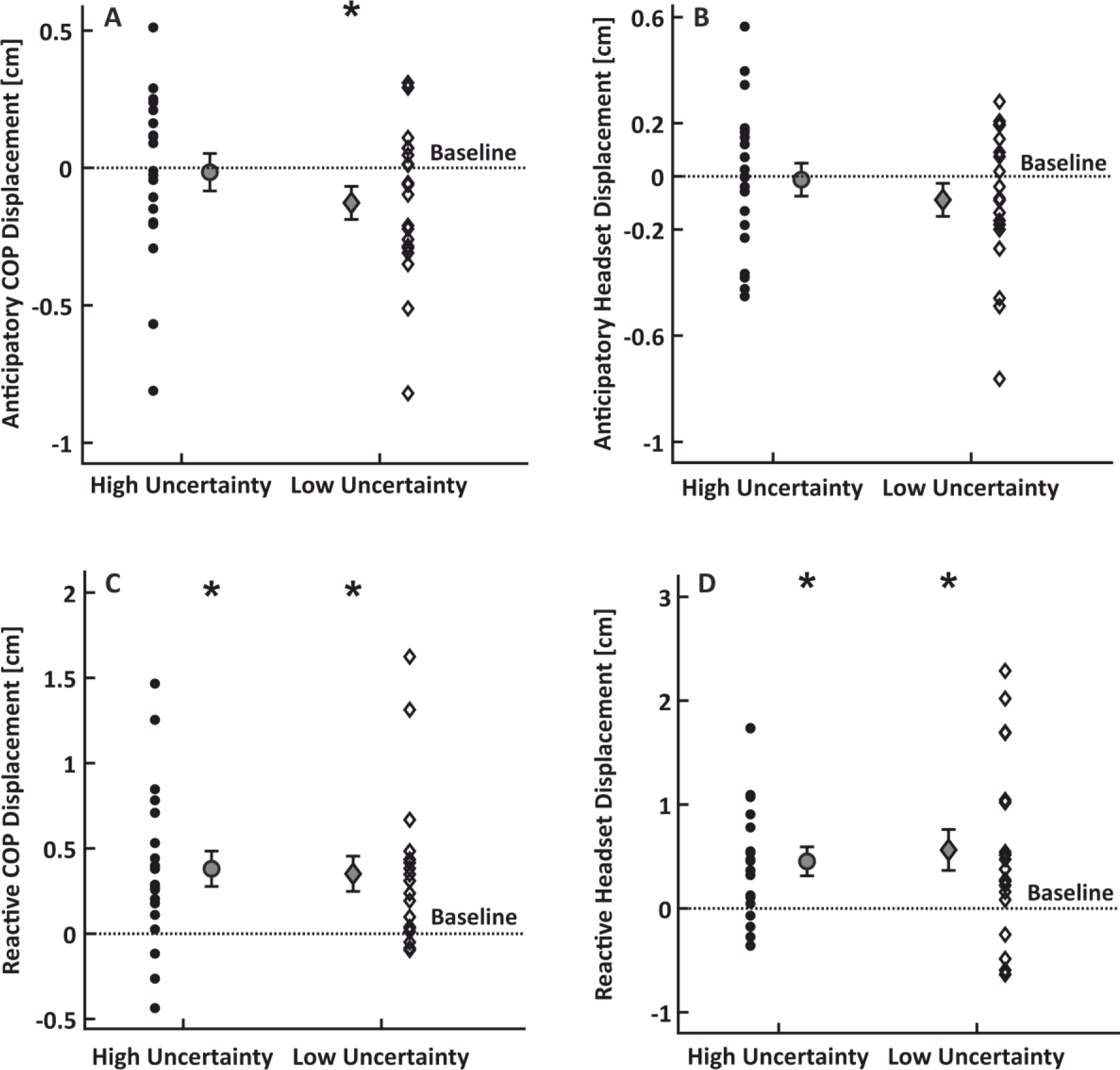
Results of the kinematic analysis. COP and head kinematic results for the anticipation and reaction periods. Panels A and B show anticipatory COP and head kinematic behavior, respectively. Panels C and D show reactive COP and head kinematic behavior, respectively. All four panels show single-subject data for the high uncertainty (circles) and low uncertainty (diamonds) conditions, as well as their corresponding means and standard errors. Dashed lines represent the baseline value and * indicates p < 0.05.

Unsurprisingly postural sway was larger when reacting to the low uncertainty perturbation relative to quiet stance, as reflected in the COP (*t_18_* = 3.402.286*, p* = 0.002*, η^2^* = 0.132) and the head kinematics (*t_19_* = 2.849*, p* = 0.005*, η^2^* = 0.097). Likewise, sway amplitudes were larger during the reaction to the high uncertainty perturbation relative to quiet stance, evident both in the COP (*t_19_* = 3.664*, p* < 0.001*, η^2^* = 0.144) and the head kinematics (*t_19_* = 3.261*, p* = 0.002*, η^2^* = 0.117). These results indicate kinematic compensations of the visual perturbation in both perturbation conditions.

#### Tactile sensitivity

We assessed tactile sensitivity by calculating tactile detection thresholds through an adaptive procedure (QUEST). Figure 2 shows the distribution of the number of trials required for the QUEST algorithm to converge, for all 173 valid detection thresholds. With an average of 8.3 ± 2.3 trials needed, the 30 trials that we presented in each condition were enough to obtain a detection threshold.

**Figure 2:**
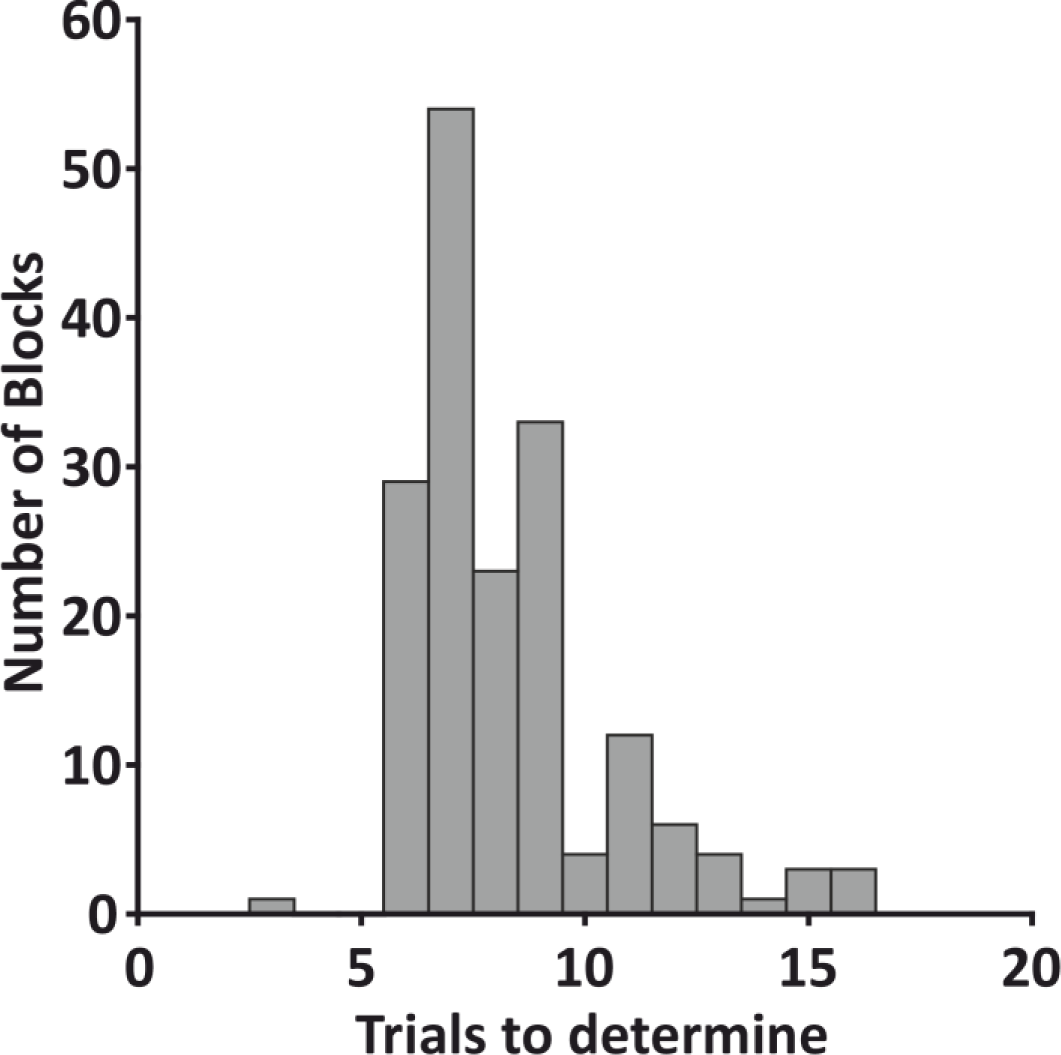
Trials until threshold estimation. Histogram of the required number of trials to determine all valid detection thresholds. The histogram shows cumulative how many trials it took to estimate the 173 detection thresholds obtained in all conditions of the experiment.

Raw detection thresholds for each time interval during the perturbation blocks are depicted in Figure 3A. Although some detection thresholds during the perturbation blocks appear to be different from those obtained during the respective standing baseline, on average the detection thresholds during perturbation blocks were not systematically different from those obtained during baseline (all *t* <= 1.933, all *p* >= 0.069, *η^2^* <= 0.047).

**Figure 3:**
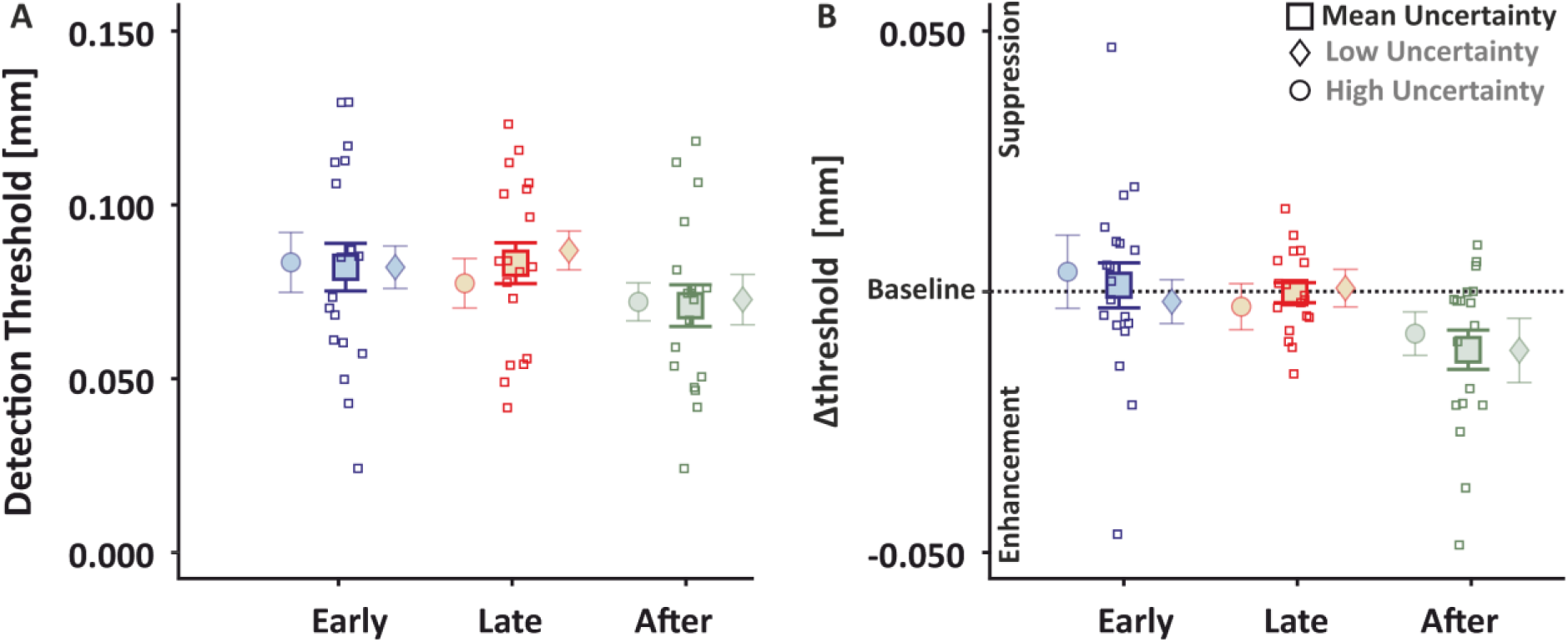
Psychophysical results Experiment 1. Detection thresholds during the perturbation blocks. Comparison of raw (A) and normalized to standing (B) detection thresholds for the three time intervals of the high (circles) and low (diamonds) uncertainty conditions are shown shaded with their average in the middle (square). Single subject data (squares) for the average are depicted. Error bars with the means and standard errors are shown.

Tactile perception during perturbation blocks relative to the standing baseline was temporally modulated as reflected in a main effect of time (*F*_2,28_ = 4.364, *p* = 0.022 *η^2^* = 0.054; Figure 3B). Post hoc t-tests revealed significantly lower detection thresholds in the “after” compared to the “early” interval (*t*_18_ = 2.942, *p* = 0.019, *η^2^*= 0.064), but similar thresholds between the “late” and “after” (*t*_18_ = 1.699, *p* = 0.201, *η^2^* = 0.022) and between the “early” and “late” intervals (*t*_18_ = 1.243, *p* = 0.224, *η^2^*= 0.012). This indicates that tactile sensitivity improved at the period after the onset of the visual perturbation, at least relative to the “early” pre-perturbation period. There was no main effect of temporal uncertainty (*F*_1,14_ = 0.364, *p* = 0.556, *η^2^* = 0.012) nor an interaction between uncertainty and time (*F*_2,28_ = 0.190, *p* = 0.828, *η^2^* = 0.004).

### Experiment 2

We conducted Experiment 2 to test if standing already causes an elevation of the tactile detection thresholds, which might marginalize any possible additional effects of feedback processing demands on tactile detection thresholds during the perturbation trials like it was previously shown for foot sole stimulation (Mildren et al., 2016). To this end, we first contrasted tactile sensitivity during quiet stance against a new sitting baseline. Indeed, standing itself led to reduced tactile sensitivity (*t*_9_ = 2.285, *p* = 0.048, *η^2^*= 0.115; Figure 4), in line with previous work (Mildren et al., 2016).

**Figure 4:**
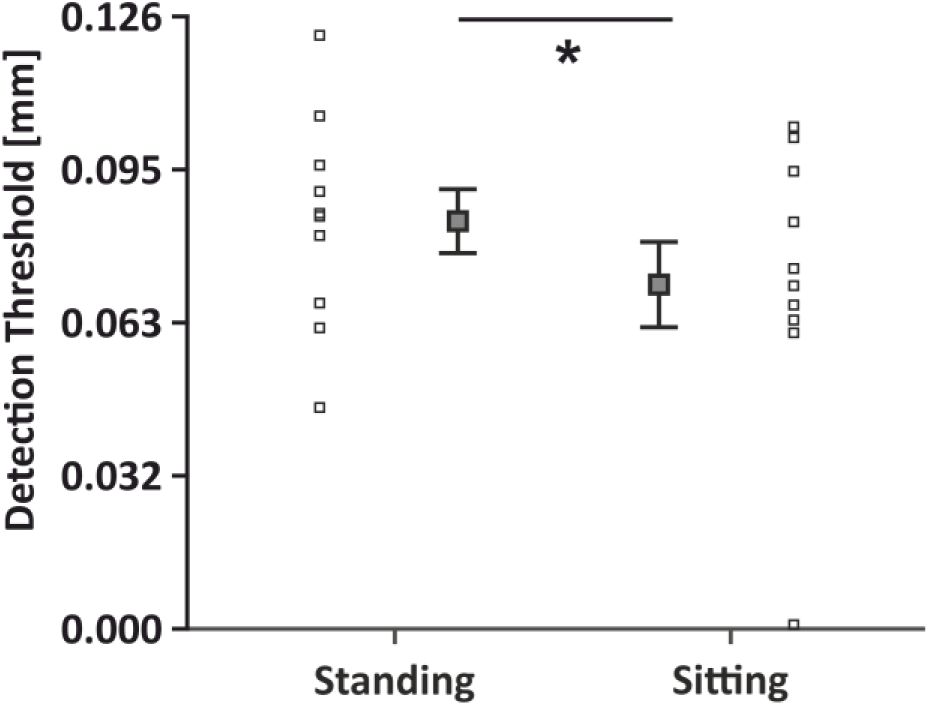
Tactile perception between standing and sitting. Comparison of raw detection thresholds between standing and sitting. This graph shows the detection thresholds for standing (left) and sitting (right) baselines. Single subject data is depicted with squares, whereas averages and standard errors are shown with squares and the respective error bars. * indicates p < 0.05.

Raw detection thresholds during the three time intervals of the perturbation blocks are shown in Figure 5A. On average these thresholds were higher than the detection thresholds obtained in the sitting baseline (all *t* >= 2.213, all *p* <= 0.027, all η^2^ >= 0.109; Figure 5B). These show a clear decline in tactile sensitivity during quiet stance around the time of visual perturbations compared to sitting.

**Figure 5:**
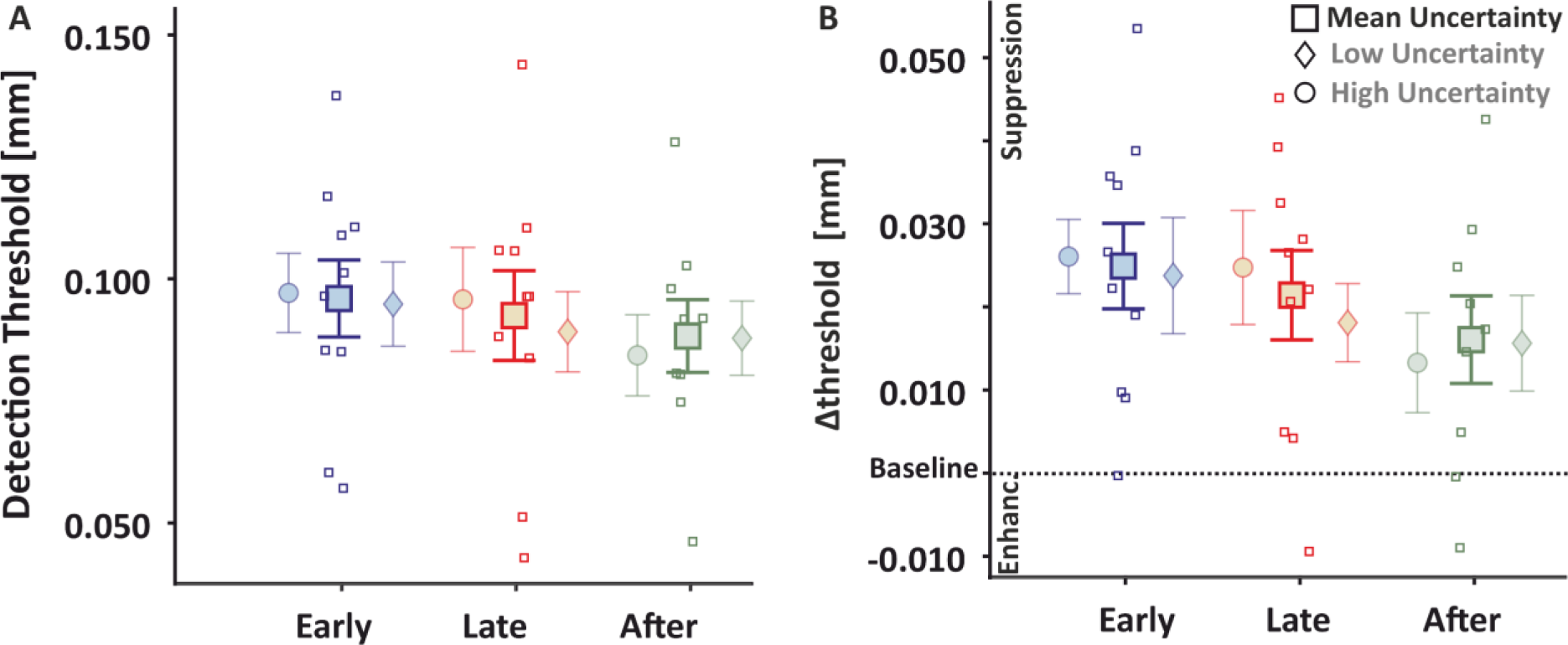
Psychophysical results Experiment 2. Detection thresholds during the perturbation blocks. Comparison of raw (A) and normalized to sitting (B) detection thresholds for the three time intervals of the high (circles) and low (diamonds) uncertainty conditions are shown shaded with their average in the middle (square). Single subject data (squares) for the average are depicted. Error bars with the means and standard errors are shown.

As in Experiment 1, the 3 x 2 ANOVA on Δthresholds revealed a main effect of time (*F_2,16_* = 16.159, *p* < 0.001, *η^2^*= 0.247; Figure 5), showing that tactile sensitivity was dynamically modulated while anticipating and reacting to perturbations. Tactile detection thresholds were reduced in the “after” compared to the “early” (*t_8_* = 5.455*, p* < 0.001*, η^2^* = 0.108) and “late” (*t_8_* = 4.113*, p* = 0.002*, η^2^* = 0.064) periods, and did not differ between the “early” and the “late” periods (*t_8_* = 1.342*, p* = 0.171*, η^2^* = 0.07). These are in line with the findings of Experiment 1 and demonstrate enhanced sensitivity around the time of perturbation, thus at moments when sensory guidance of posture gains importance. There was again no main effect of temporal uncertainty (*F_1,8_* = 1.089, *p* = 0.327, *η^2^* = 0.037), nor an interaction (*F_2,16_* = 0.642, *p* = 0.539, *η^2^* = 0.024).

## Discussion

We examined whether tactile sensitivity on the lower limb is modulated by feedback processing demands. To this end, we used an adapted version of the classical moving room paradigm and examined the temporal modulation of tactile sensitivity on the lower leg during standing, while preparing for and reacting to visual perturbations that occurred with low or high temporal uncertainty. We measured tactile sensitivity at the leg at different times during this task. We show that feedback processing demands modulate tactile sensitivity leading to enhanced sensitivity after the onset of the perturbation, i.e. at the moment close to the onset of the reactive postural response. This demonstrates that tactile sensitivity on the standing leg during balance retention is dynamically tuned to the dynamics of the perturbation and the need to sample sensory feedback from the probed body part.

The increased tactile sensitivity on the lower leg after the onset of the perturbation indicates an upweighting of somatosensory feedback signals from the standing limb around that moment. This is inconsistent with previous findings that found *reduced* tactile sensitivity right before and during single finger movements (Chapman et al., 1987; Williams et al., 1998), reaching and grasping actions (Colino & Binsted, 2016) or foot extensions (Morita et al., 1998). This apparent discrepancy can be explained by differences in the task relevancy of the sensory information from the probed limb. In our study, somatosensory information from the lower leg is important to control posture, particularly around the onset of the reactive response. This explanation is in line with findings showing increased somatosensory-evoked potentials when stimulating the foot sole shortly before initiating a step (Mouchnino et al., 2015). Our results expand the set of studies showing that tactile sensitivity can be modulated during an action (e.g., Colino & Binsted, 2016; Voudouris et al., 2019; Voudouris & Fiehler, 2022; Mouchnino et al., 2015; Voudouris & Fiehler, 2021), especially when feedback signals from the probed limb are important for the task, suggesting that tactile modulation around human movement is a dynamic process.

We also examined the effect of temporal uncertainty related to the moment when the visual perturbation was triggered. There is evidence that sensorimotor uncertainty can influence the strength of tactile sensitivity (Wolpe et al., 2016; Klever et al., 2019; Voudouris et al., 2019). Here we explored if low temporal uncertainty about the onset of the perturbation would lead to stronger reliance on predictive control when task demands increase. Although we found anticipatory motor adjustments before perturbations of low temporal uncertainty, we have no evidence that tactile sensitivity was influenced as well. However, we cannot exclude the possibility that the uncertainty about the onset of the perturbation was too small to impact tactile perception since the general trial structure of the two perturbation conditions was comparable.

Interestingly, in Experiment 1, tactile sensitivity during standing was not different between perturbation and quiet stance. This indicates that standing upright already leads to increased tactile detection thresholds. Indeed, when compared to sitting in Experiment 2, tactile sensitivity during standing was significantly reduced, in line with previous findings that assessed tacile sensitivity on the foot sole (Mildren et al., 2016). Tactile suppression during upright stance may stem from one or more sources. One possibility is that changes in skin properties due to muscle activation or leg configuration have altered the mechanical properties of the skin around the probed area and thereby worsend tactile detection (Provancher & Sylvester, 2009). It is also possible that backward masking from the activation of the lower leg muscles increased sensory noise and thus made the brief vibrotactile probes less conspicuous. Indeed, standing itself is accompanied by a sinusoidal movement (Winter, 1995) and postural adjustments (Warnica et al., 2014). When the stance is less stable e.g. the base of support is reduced, the activity of the lower leg muscles increases (Amiridis et al., 2003) and this increase in peripheral activity may mask the probing tactile stimuli (Williams & Chapman, 2002; Chapman & Beauchamp, 2006). However, so far there is no clear evidence that increased force production and hence muscular activity per se can modulate tactile suppression (Broda et al., 2020).

The sitting baseline occurred always at the end of the experimental procedure. This might render the reduced tactile sensitivity during standing as a by-product of improved tactile detection performance at the later (sitting) baseline simply due to practice. This could lead to lower detection thresholds toward the end of the experiment compared to the preceding standing tasks. This is unlikely in our experiments for two reasons. First, the standing baseline tasks that were presented at the end of Experiment 1 did not yield lower detection thresholds than the earlier presented perturbation tasks. Thus, the lower detection thresholds during sitting, which was the last block in Experiment 2, are unlikely to arise simply because of a longer exposure to the tactile task. Second, tactile detection thresholds, at least on the hand, do not differ between sessions separated by up to 30 minutes (Voudouris & Fiehler, 2022). All in all, we have no evidence that the reported effects may be caused by a possible improvement in tactile detection over the course of the experiment.

Our kinematic results demonstrate systematic anticipatory adjustments prior to the perturbations when temporal uncertainty was low. Unsurprisingly, there were no anticipatory adjustments prior to the perturbation of high temporal uncertainty. This is in line with previous work showing anticipatory postural adjustments prior to predictable perturbations (Aruin & Latash, 1995; Eikema et al., 2013). The anticipatory effects were mostly evident in the COP, while the head kinematics followed a similar but more variable pattern. This might be due to the sensorimotor system prioritizing the stabilization of the head over the trunk, as head stabilization is a common biological strategy. Stabilizing the head could reduce transformations of the sensory signals triggered during head movements or could provide a stable reference frame for the task at hand (Berthoz & Pozzo, 1994). We also confirmed that participants react to new visual information by compensatory COP and head adjustments evident after the onset of a perturbation, and independently of its temporal uncertainty. The reactive amplitudes may appear rather small, but it is important to note that our participants had a narrow base of support (i.e. feet close together), which would make it disadvantageous to exert large sways that could shift the center of pressure outside this narrow base of support and lead to postural instability. Indeed, when the base of support induces greater postural instability, body reactions have much smaller amplitudes (Voudouris et al., 2013).

Together, our results demonstrate temporal modulation of tactile sensitivity on the lower limb when maintaining whole-body balance around visual perturbations. The temporal tuning of tactile sensitivity when anticipating and reacting to a visual perturbation provides evidence for the idea that tactile sensitivity is modulated dynamically to the feedback demands. This upweighting of sensory feedback signals can lead to improved tactile sensitivity on a moving body part when somatosensory feedback signals from that part gain relevance for the sensory guidance of the movement.

## Limitations of this study

Even though there were anticipatory postural adjustments before the low uncertainty perturbation and no adjustments before the high uncertainty perturbations, we did not detect differences in tactile sensitivity between these two conditions. We assume that the uncertainty was large enough to cause small postural adjustments but too low to elicit measurable changes in tactile sensitivity. Another limitation refers to the origin of the tactile modulation. While we demonstrate that tactile sensitivity is dynamically modulated in both experiments, our experimental paradigm cannot pinpoint the exact underlying mechanisms of this phenomenon. We describe possible explanations of this phenomenon in the Discussion.

Due to the low spatial specificity in the modulation of tactile perception (Voudouris & Fiehler, 2017b) we were able to probe tactile perception on the lower leg. It is possible that the observed effect on increased sensitivity just before reactive adjustments would be even stronger and an anticipatory increase in tactile sensitivity observable when probed at the foot sole as done in other studies (Mouchnino et al., 2015). However, our device did not allow for such testing opening an interesting avenue for future research.

## Author contributions

**Conceptualization** F.W., K.F., D.V.

**Methodology** F.W., K.F., D.V.

**Software** F.W.

**Validation** F.W., K.F., D.V.

**Formal Analysis** F.W.

**Investigation** F.W.

**Resources** F.W., K.F., D.V.

**Data Curation** F.W.

**Writing – Original Draft** F.W.

**Writing – Review & Editing** F.W., K.F., D.V.

**Visualization** F.W.

**Supervision** K.F., D.V.

**Project Administration** F.W., K.F., D.V.

**Funding Acquisition** K.F, D.V.

## STAR★METHODS

### Key Resources Table

| Resource | Source | Identifier |
| --- | --- | --- |
| *Software and Algorithms* |  |  |
| Matlab 2021a | MathWorks | https://de.mathworks.com/products/matlab.html |
| QUEST | Psychtoolbox 3.0.18 | http://psychtoolbox.org/ |
| JASP 0.17.1 | JASP | https://jasp-stats.org/ |
| Unity 2021.3.4f1 | UNITY | https://unity.com/ |
| *Hardware* |  |  |
| Tactor | Engineering Acoustic Inc. | https://eaiinfo.com/ |
| Forceplate | Advanced Mechanical Technology Inc. | https://www.amti.biz/ |
| VIVE Pro Eye | HTC Corp | https://www.vive.com/de/ |

### Resource availability

#### Lead contact

Further information and requests for resources should be directed to and will be fulfilled by the lead contact, Fabian Dominik Wachsmann

#### Data and Code availability

Behavioral and psychophysical data are publicly available after acceptance at https://osf.io/jdzwa/. JASP/R-Code for statistical analysis will be attached as .txt file.

### Method details

#### Experiment 1

##### Participants and apparatus

We recruited 20 participants (24.4 ± 4.8 years old; range: 19-39, 11 ♀ 9 **♂**) for a one-session virtual reality experiment. One additional participant was recruited but did not finish data collection due to circulatory problems. Participants were free from any known neurological or musculoskeletal issues at the moment of the experiment and had normal or corrected-to-normal vision. Upon their arrival, participants signed an informed consent approved by the local ethics committee of the Justus Liebig University Giessen in accordance with the “World Medical Association Declaration of Helsinki” (2013, except for §35, pre-registration). At the end of the experiment, participants received for their efforts either 8€/hour or course credits.

A custom-made vibrotactile stimulation device (Engineer Acoustics Inc., Florida, US), which will be referred to as a “tactor”, was attached to the *musculus gastrocnemius lateralis* of the participant’s right calf. This tactor was fixed at 3 cm medial of 2/3 of an imaginary line connecting the heel with the *caput fibulae.* Thus, the tactile device was fixed on the belly of the muscle that was expected to mainly control the retention of upright stance in our experiment. Care was taken that loose wires or clothing could not be mistaken as vibrations and that they would not hinder the performance of the task. Participants stood barefoot or with their socks on having their feet closed together on a force plate (AMTI, Massachusetts, US) that sampled the center of pressure (COP) at 300 Hz. They also wore a head-mounted display (HTC Vive Pro Eye, Taiwan) that presented a virtual environment and sampled positional data at a rate of 90 Hz. Participants were then immersed in a 3 x 3 x 6 m (width, height, depth) room with black and white striped side walls and a grey front wall. At 1.85 m above the floor, a circular fixation point with a 60 cm radius was presented (Figure 6). The virtual environment was a custom-made Unity version 2021.3.4f1 (Unity Technologies, San Francisco, US) build.

**Figure 6:**
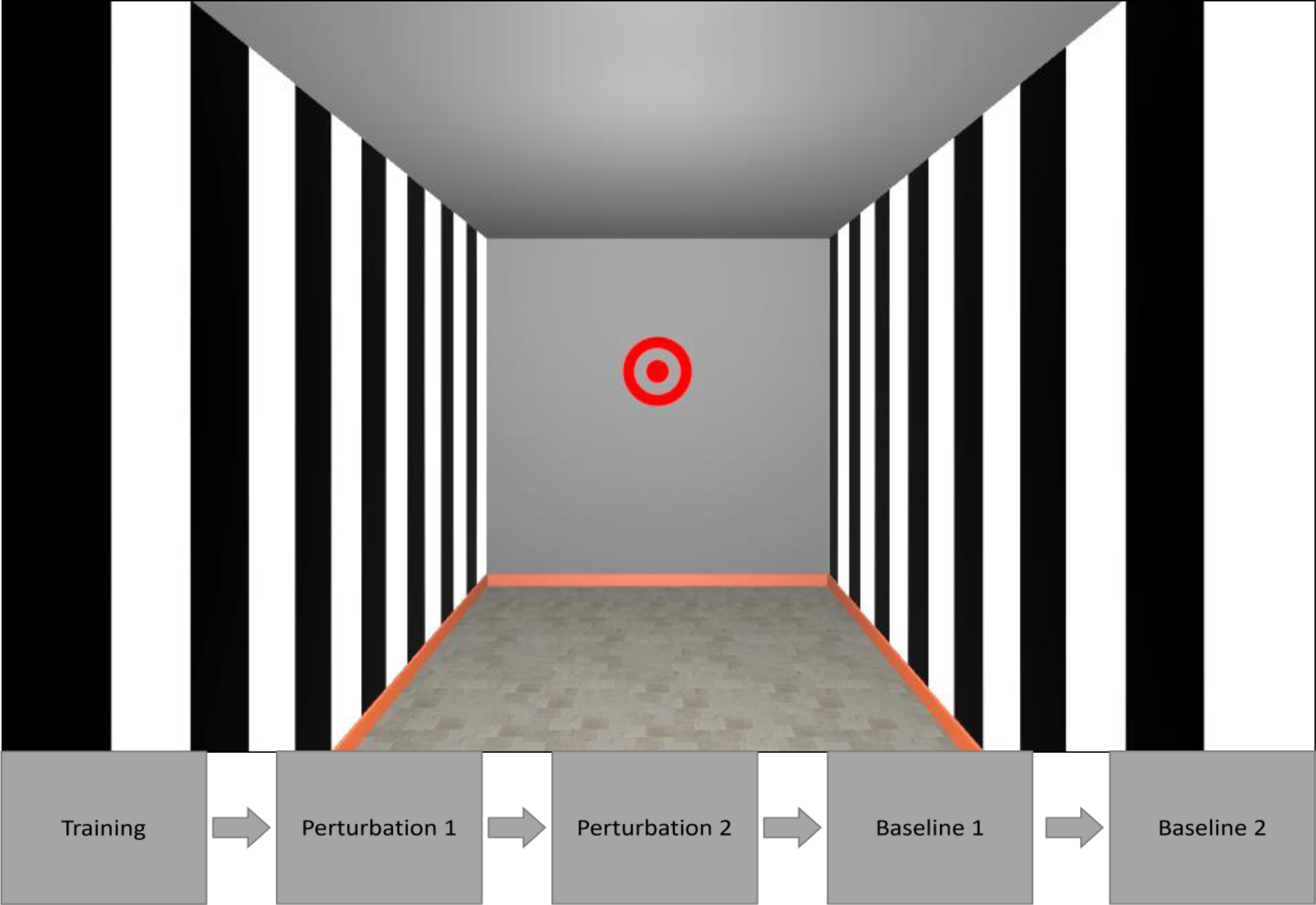
Virtual reality setup. Picture of the virtual environment as seen from the participants. Depiction of the virtual room, with the red circular fixation point appearing at the front wall. The lower panel shows the block order.

##### Procedure

The experiment consisted of five separate blocks of trials, with each trial lasting 7 s. The first block was always a set of training trials. Here, a countdown (3, 2, 1) appeared above the fixation point on the front wall, with each digit being presented for 0.75 s. At the end of this countdown, a visual perturbation occurred, which consisted of the complete room moving in such a way that the front wall approached the participant with a velocity of 3 m/s for 5 m. Thus, the movement of the room toward the participant lasted 1.67 s. After each trial, the room instantly recovered back to its original configuration. After this training block, two perturbation conditions were presented in a random order across participants. In each of these two blocks, the room could be perturbed at a moment of low or high temporal uncertainty. In the low uncertainty condition, trials started with a fixed delay of 0.75 s, followed by a 2.25 s period of countdown, and the visual perturbation occurred immediately after that countdown. In other words, the perturbation always occurred 3 s after the onset of the trial. In the high uncertainty condition, a trial would start without a countdown, and the perturbation would occur within a random moment between 2.25 and 3 s from the beginning of the trial. The absence of the countdown and the 0.75 s random delay increased the uncertainty about the exact moment of the perturbation onset. These resulted in a difference in the uncertainty of when the perturbation would occur between these two conditions, while also allowing us to capture the same time window during which the countdown appeared, and thus we could examine possible anticipatory sensorimotor responses prior the perturbation in both perturbation conditions. Upon completion of these two perturbation conditions, two blocks followed where we assessed baseline sensory and motor performance: one block with and another block without a visual countdown, but importantly both blocks without any visual perturbation. In other words, these two baseline conditions were identical to the two perturbation conditions but now the virtual room never moved. The order of the two baseline conditions was identical to the order of the perturbation conditions, and this order was randomized across participants. Participants were explicitly informed prior to the onset of each block about the condition that they would be exposed to. In all blocks, participants were asked to fixate the fixation point at the center of the front wall all times.

To probe tactile sensitivity, we delivered a brief (50 ms, 250 Hz) vibrotactile stimulus of varying intensities at the participants’ right calf muscle at various moments during each trial, and participants had to report whether they felt a vibration or not. This tactile stimulus is not relevant for the postural task, but is used as a proxy to assess tactile sensitivity; a procedure commonly applied in previous work (e.g., Buckingham et al., 2010; Colino & Binsted, 2016; Voudouris & Fiehler, 2021). In the two perturbation conditions, the probing tactile stimulus was delivered at a random moment during one of three possible intervals relative to perturbation onset: two at the pre-perturbation period (“early”: −2.25 to −1.50 s, and “late”: −1.50 to −0.75 s) and one at the post-perturbation period (“after”: 0.00 to 0.75 s; Figure 7). We presented 30 trials at each of these three intervals in a randomized order, for a total of 90 trials per perturbation block. In each of the training and two baseline conditions there was a single time interval during which the tactile probe could be presented: for the training and baseline condition with the countdown the stimulus was presented between 0.75 – 3.75 s, whereas for the baseline condition without the countdown it was presented between 0 – 3.75 s, always relative to the onset of the trial.

**Figure 7:**
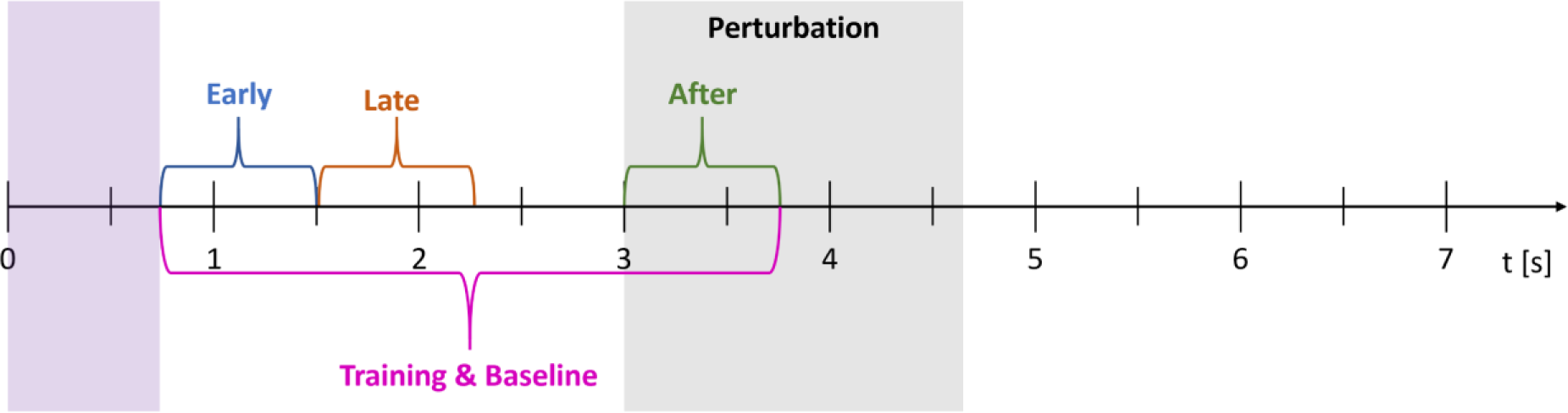
Timecourse of experimental trials. Schematic timeline of a single trial of the perturbation condition. The probe tactile stimulus in the two perturbation blocks could be presented at one of three possible intervals: “early” (blue), “late” (brown), and “after” (green), whereas in the baseline and training conditions, it could be presented at any moment during a single interval (purple). The onset of the perturbation is depicted here as starting at the 3^rd^ second of the trial, but this could vary in the condition with the high uncertainty perturbation. The duration of the perturbation is shown with the grey area.

The intensity of vibration in each trial was determined using a QUEST algorithm (Psychtoolbox Version 3.0.18). As a Bayesian approach, the QUEST algorithm uses prior knowledge to estimate the parameters of a psychometric function. As the model function we used a Weibull distribution because it can represent a wide range of response patterns. We chose a β of 3.5 for the distribution which is the suggested value for 2AFC tasks in the QUEST algorithm (Watson & Pelli, 1983). The prior mean and standard deviation for the training block were estimated from a pilot study where participants just stood with their feet closed (N=16, exposed to 50 stimuli of varying intensities, presented at the calf muscle of the right leg). We used the mean of the estimated probability density function as probe intensity for the next trial. For all the other conditions we used the estimated threshold of the training block as prior mean (Watson & Pelli, 1983; May & Solomon, 2013).

In each trial, participants were instructed to verbally report whether or not they felt a vibration. They were instructed to immediately respond once they felt a vibration to avoid memory effects, but they were also explicitly asked to respond after the trial ended if no answer was meanwhile given. These verbal responses were then registered by the experimenter via button presses to a computer, which induced a short delay (0.5 – 1 s) between trials. To prevent fatigue, participants could have a break after every trial on request. Breaks of ∼1 minute were enforced every 30 trials and longer breaks of ∼5 minutes were introduced after 90 trials. During these longer breaks, participants were required to sit and were allowed to remove the headset. This resulted in a duration of around 20 minutes for each perturbation block, with a total experimental time of 90 minutes. Trial control, data acquisition, and data analysis were handled by custom-made Matlab R2021a software (The Mathworks, Massachusetts, US).

##### Data analysis

The COP and headset kinematics were analyzed only in the anterior-posterior (negative to positive values, respectively) direction as the visual perturbation happened in this direction. In a first step, we excluded trials from all kinematic and psychophysical analyses with data that were continuously missing for at least 100 ms (1.2%; only COP data were ever missing). Afterward, we normalized every sample of each trial to the average position of the first 0.5 s of that trial, which we consider as a period of quiet stance because there was nothing happening to the room. The resulting normalized data was corrected for outlier data points by replacing values with a displacement larger than 15 cm with no value (this only affected COP data), and then smoothed utilizing a symmetric moving average with a window size of 100 ms.

Anticipatory kinematic adjustments were quantified as the maximum absolute COP and head amplitude during the last 2.25 seconds prior to perturbation onset. To quantify reactive kinematic behavior we first determined the maximum positive (posterior) COP and head displacement during the first 3.5 seconds following the onset of the perturbation. This corresponds to about 2 times the perturbation duration. We calculated both components for every trial of each subject and averaged across all trials for each uncertainty level. Determining the maximum positive COP and head position values to quantify motor responses was important because any reaction to the perturbation would result in the body leaning backward and thus to a positive COP and head position value. We then quantified reactive kinematic adjustments of the COP and head as the difference between their maximum value and their respective minimum that occurred between the perturbation onset and that maximum (see Figure 8). Both of the previous measures were normalized with respect to the values calculated from their respective baseline periods (quiet stance) to account for individual differences in standing. Thus, values greater than zero indicate larger displacements during the perturbation blocks relative to the respective baseline.

**Figure 8:**
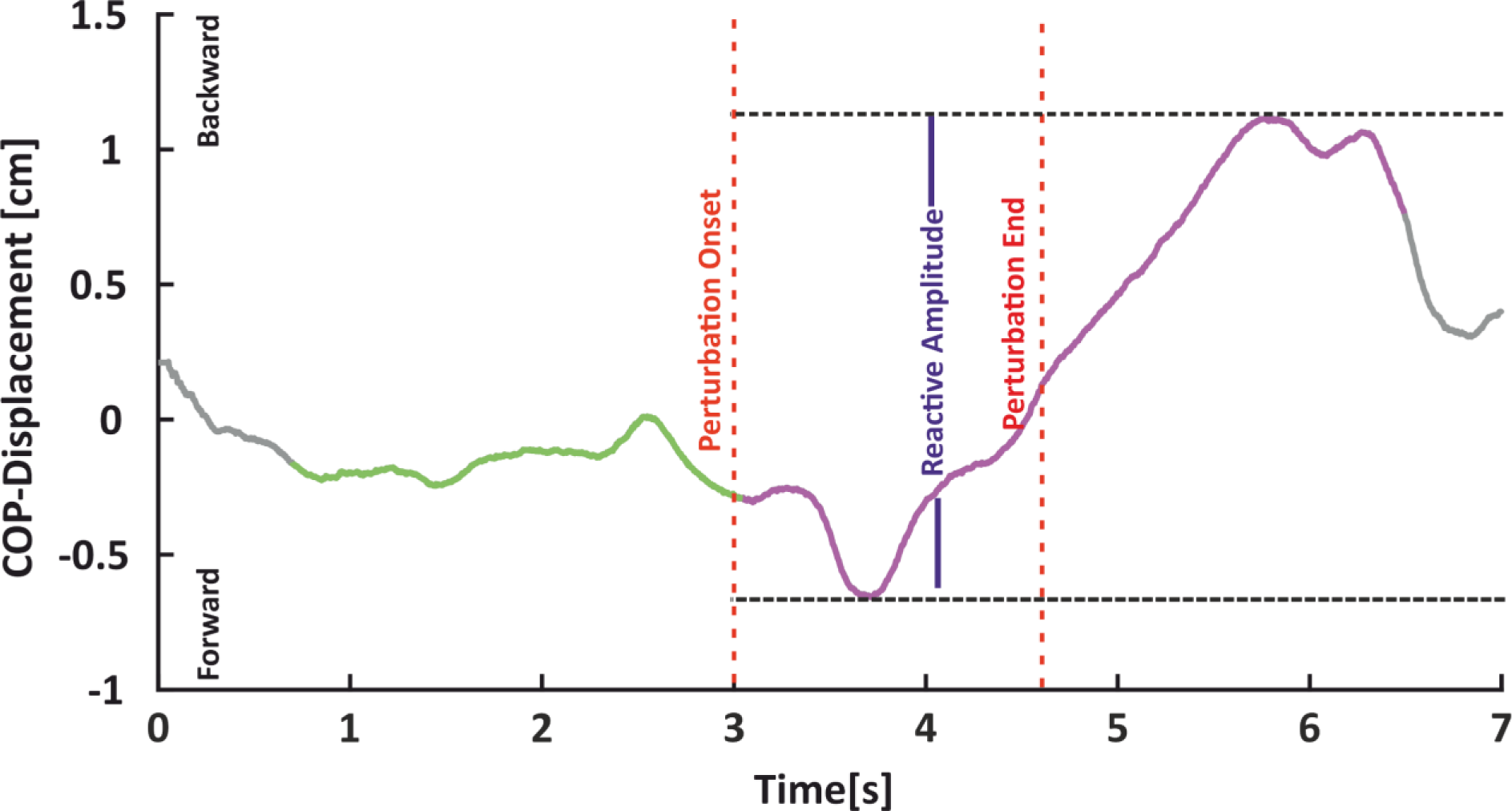
Exemplary kinematic data. Temporal profile of the center of pressure (COP) time course in the antero-posterior direction of a low uncertainty trial. The time window of perturbation (red dashed lines), the anticipatory period (green path), the reactive period (purple path), and the reactive amplitude (blue solid line) are depicted.

We determined tactile detection thresholds for each condition and participant. This was quantified as the first trial after which the probe stimulus intensity recommended by the QUEST did no longer change value. It is important to note that our device accepts only integer values as inputs for the tactile stimuli, but the values recommended by the QUEST could also include decimals. In these cases, we always rounded the suggested value to the closest integer. Thus, although the QUEST suggested slightly different intensities in the trials after the detection threshold was determined, due to rounding we presented the same stimulus intensities. We expected some fluctuation in the responses to those trials due to natural randomness in detecting stimuli around one’s own detection threshold. For this reason, we considered detection thresholds as invalid if participants gave the same (detected or not-detected) response in the remaining trials after the detection threshold was determined. Those thresholds (4.2%) were excluded from further analysis and their corresponding kinematic data. To quantify tactile modulation during perturbation relative to quiet stance trials while also accounting for individual differences in tactile sensitivity, we subtracted each participant’s baseline detection threshold from their threshold obtained at each time interval of the respective perturbation block (Δthreshold), a procedure in line with previous work (Fuehrer et al., 2022). To closer align the kinematic with the perceptual results we included in the kinematic analyses of each condition and participant only those trials that were used to estimate the associated detection thresholds. Kinematic data from trials where the estimated threshold was judged invalid was omitted.

##### Statistical analysis

We first examined whether anticipatory behavior was evident prior to perturbation compared to baseline by testing if the maximum displacement in COP and head kinematics during the anticipatory period deviated from the respective values during quiet stance. Because we subtracted the maximum displacement in baseline from the respective value in the perturbation blocks, we quantified possible effects by using one-sided one-sample t-tests against zero for the low uncertainty (based on the directed hypothesis) and two-sided t-tests for the high uncertainty condition (based on the undirected hypothesis). Similarly, we tested whether the reactive component in each perturbation condition deviated from zero (i.e. baseline) using one-sided one-sample t-tests.

To first explore possible modulation of tactile sensitivity during the perturbation blocks relative to baseline, we submitted the obtained Δthreshold values to six separate two-sided one-sample t-tests against zero (i.e. baseline). Our main interest was whether tactile perception would be temporally tuned while anticipating and reacting to a perturbation. We also explored whether there would be any effects of the temporal uncertainty of the perturbation. To test for possible effects of the time interval of tactile probing (“early”, “late”, “after”) and the type of perturbation (low vs high uncertainty) on tactile sensitivity, we used a 3×2 repeated measure ANOVA. Significant main effects were explored with post-hoc t-tests, corrected for multiple comparisons wherever necessary using the Holm procedure. All p-values were Holm-corrected and all t-values were reported as absolute values. The type one error threshold was set to 0.05 and effect sizes are reported as *η^2^* following the calculations and recommendations from Correll et al. (2020). Statistical analyses were conducted in JASP version 0.17.1 (University of Amsterdam, Netherlands). Datapoints in any of the 6 experimental or 2 baseline conditions that were outside of the 3.5 interquartile range were excluded from the statistical analysis of that condition as outliers (< 1% for kinematic and 0% for psychophysical data). If any baseline value was excluded, the three values in the associated perturbation condition were also excluded since normalization was impossible.

#### Experiment 2

In Experiment 1 we examine the temporal modulation of tactile sensitivity when coping with visual perturbations of high or low temporal uncertainty during upright stance. To this end, we compare tactile perception when standing and being confronted with a perturbation to when standing without having to cope with any perturbation. However, standing itself may already influence tactile sensitivity on the leg, for instance by masking feedback signals, or by changes in skin and muscle-related properties on the probed body part. To account for such possibilities, we conducted Experiment 2, where we examined whether tactile detection thresholds at the lower leg differ between quiet stance and sitting. We further examined whether tactile modulation followed similar patterns as those observed in Experiment 1.

##### Participants and experimental design

We recruited 10 new participants (25.3 ± 4.0 years old; range: 21-32; 9 ♀ 1 **♂**) for a one-session experiment, while two additional participants did not finish data collection due to circulatory problems or incompliance with task instructions. The procedure was almost identical to Experiment 1, but now participants performed six blocks of trials: a training block, two perturbation blocks, two standing baselines, and after these, an additional sitting baseline. During this sitting baseline, participants wore the head-mounted display and sat relaxed on a chair with their knees flexed at ∼90° and both feet touching the floor. In this sitting baseline, participants performed 30 trials that were constructed identically as in the other two standing baseline blocks. In all six blocks of Experiment 2, participants saw the same room as in Experiment 1. The experimental procedure and analyses were identical to those reported for Experiment 1, except for the details mentioned below.

##### Data and statistical analysis

As we were interested in the modulation of tactile sensitivity by standing, we focused our analysis on tactile detection thresholds. In contrast to Experiment 1, detection thresholds that were obtained during the two perturbation blocks of Experiment 2 were now normalized with respect to the new sitting baseline, resulting again in six Δthreshold values per participant. The analysis of the participants’ responses after the detection threshold resulted in excluding 1.5% of the total detection thresholds and their corresponding kinematic data. Furthermore, and since we did not observe any differences in the two standing baselines in Experiment 1 (*t*_19_ = 0.621, *p* = 0.542, *η^2^* = 0.005) or Experiment 2 (*t*_9_ = 0.263, *p* = 0.798, *η^2^* = 0.002), we now averaged across the two standing baselines thresholds for each participant.

A two-sided paired t-test was performed to test for differences between the standing and the sitting baselines. To further examine whether tactile detection thresholds increase during the perturbation blocks relative to the sitting baseline, we contrasted the six Δthreshold values against zero (i.e. baseline) with six separate one-sided one-sample t-tests. Additionally, we submitted the Δthreshold values to a 3×2 repeated measure ANOVA to investigate for main effects of the time of probing (“early”, “late”, “after”) and of type of perturbation (low vs high uncertainty) on tactile modulation, as in Experiment 1.

